# RAEM: random-access electron microscopy for revisitable 3D imaging

**DOI:** 10.64898/2026.06.18.732873

**Authors:** Ishaan Singh Chandok, Milan Patel, Yuelong Wu, Daniel Berger, Richard Schalek, Jeff W. Lichtman, Aravinthan D.T. Samuel, Yaron Meirovitch

**Author notes:** Correspondence (ISC); (JL); (AS); (YM).

## Abstract

Volume electron microscopy is essential for understanding cells, tissues, and neural circuits in their native 3D context, but many biological specimens are too large to image exhaustively at nanometer resolution. Researchers therefore must choose between broad anatomical context and ultrastructural detail. We introduce random-access electron microscopy (RAEM), a framework for studying fixed tissue repeatedly across scales rather than imaging it once at a single resolution. RAEM first builds a lower resolution 3D survey of the specimen, then uses accumulated human or AI-derived knowledge of that volume to guide the microscope back to selected physical sites for high resolution imaging. By linking reconstructed 3D coordinates to precise electron-beam positions on the original sections, RAEM enables targeted imaging of membranes, vesicles, and other nanoscale structures within specimens that would be impractical to image exhaustively. We demonstrate RAEM with vesicle-resolved imaging of synaptic boutons in human cortex, targeted imaging of more than one million human cortical mitochondria, hierarchical imaging of a nematode nervous system, and retrospective targeting of a previously published petabyte-scale human cortical volume. RAEM turns serial-section EM into a query-driven, multi-resolution approach for scalable biomedical discovery.

## Introduction

Traditional serial-section electron microscopy (ssEM) pipelines treat biological specimens uniformly. Sections are typically imaged in a single pass, with every pixel acquired at the spatial resolution (pixel size and dwell time) required for the most difficult-to-resolve ultrastructural features in the sample. Yet many EM studies require such imaging only for a small fraction of the total specimen volume (**Figs. 1a, 1b**). Large portions of the specimen can instead be omitted or acquired at lower resolution without compromising the reconstruction. As a result, enormous imaging effort is expended on complete volumes, even though structurally complex features, such as synapses, might be sparse and irregularly distributed.

**Figure 1.**
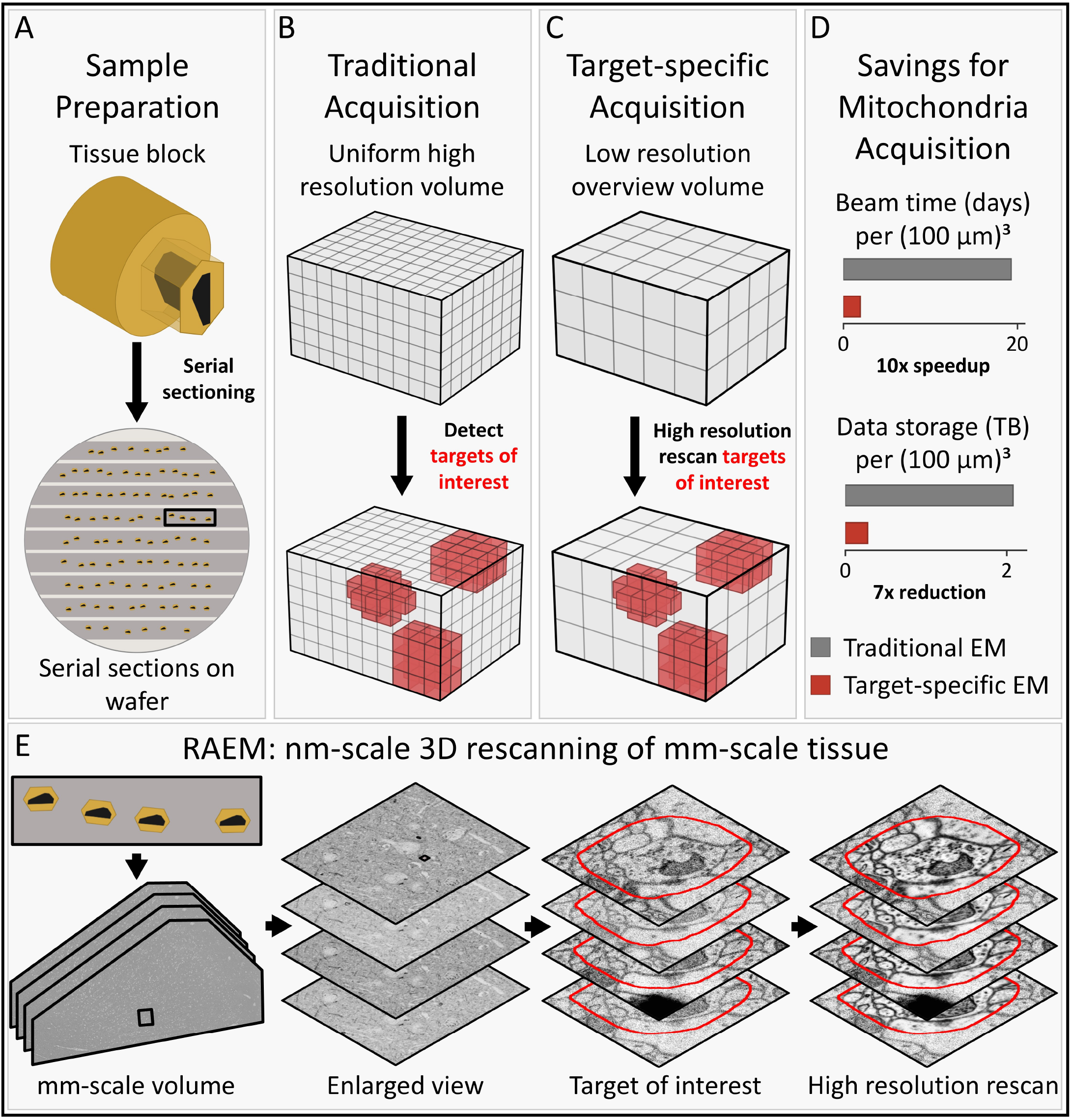
Overview of the target-specific EM workflow and the RAEM pipeline. **A**. The typical ssEM pipeline begins with sample preparation, where a tissue block is sectioned into serial sections mounted on a substrate such as tape on a wafer. **B**. In traditional EM acquisition, a sample is imaged everywhere at a single high resolution (small pixel size and long dwell time), and only targets of interest (in red) are analyzed while targets of non-interest are excluded from subsequent study. **C**. In a target-specific EM workflow, an overview volume is first acquired, targets of interest are detected, and these targets are then revisited for targeted higher resolution imaging. Only these targets are imaged at high resolution. **D**. Estimated beam time and data storage needs for imaging all mitochondria at high resolution in a (100 *µm*)^3^ volume of human cortex, comparing traditional and target-specific EM. **E**. The RAEM pipeline enables arbitrary target-specific acquisition by performing nanometer-scale 3D targeted imaging across millimeter-scale tissue.

To date, large-scale connectomic reconstructions have depended on conventional ssEM workflows. In early brain-mapping efforts, comprehensive and unbiased tissue reconstruction was necessary. Efficient acquisition of these datasets required high-throughput, specialized, and costly platforms, including the Zeiss 61- and 91-beam scanning electron microscopes and TEMCA2, which remain accessible to only a few laboratories ^**1, 2**^.

The next generation of hypothesis-driven studies, however, does not require comprehensive reconstruction of entire sample volumes. Instead, these applications call for targeted imaging of specific cells, subcellular compartments, or ultrastructural features sparsely embedded within much larger tissue volumes. Examples include quantifying changes in synapse morphology across physiological states ^**3**^, determining how organelle ultra-structure is altered in disease models ^**4**^, and measuring remodeling of synaptic architecture following plasticity induction ^**5, 6**^. The time and cost required to pursue such questions at scale using traditional ssEM workflows have been prohibitive.

Another route to scale is to make imaging selective rather than uniform. Target-specific ssEM can be achieved by separating rapid acquisition of a 3D overview from high resolution acquisition of selected targets (**Fig. 1c**). The overview provides the volumetric context needed to identify regions of interest, which may be small and sparsely distributed throughout the specimen and must then be physically revisited for high resolution imaging. Importantly, both steps can in principle be performed using widely available single-beam SEM hardware.

In earlier work, we developed SmartEM, an AI-based approach that improved imaging efficiency within a single microscope field of view on a single-beam SEM ^**7**^. SmartEM acquired a rapid 2D overview at low resolution and then rescanned only those pixels whose high resolution acquisition quantitatively improved segmentation of biological structures.

Here we develop a new mode of image acquisition that uses 3D context to reconstruct ultrastructural targets distributed across multiple serial sections. For example, imaging all mitochondria at high resolution within a (100 *µ*m)^3^ volume of human cortex would require approximately 19 days of beam time and 2 TB of storage using uniform acquisition at 4 nm and 800 ns on a single-beam SEM. By contrast, a workflow that first acquires a 3D overview at 16 nm and 400 ns and then images only detected mitochondria, occupying approximately 7% of each serial section area, would require approximately 2 days and 0.3 TB. This corresponds to an approximately tenfold reduction in beam time and a sevenfold reduction in storage (**Fig. 1d**).

Applying global 3D context in a target-specific workflow poses substantial practical challenges. The microscope must return to target locations on sections mounted on substrates spanning tens of centimeters while imaging with nanometer resolution and nanometer-scale accuracy, thereby bridging eight orders of magnitude in scale. Mapping a target in the digital volume back to its physical location on the substrate is limited by stage-motion imprecision, sample drift, deformation, and alignment error. These errors must therefore be corrected in real time to enable accurate targeted imaging and seamless fusion of the acquired data into the original 3D overview.

We introduce the random-access electron microscope (RAEM), a framework that uses ssEM to generate a 3D digital overview of a biological sample and then selectively acquire desired targets with nanometer-scale accuracy and resolution. In RAEM, the overview serves as a digital index of the specimen: targets identified within this volume can be revisited physically on the sample, imaged at higher resolution, and fused into the same coordinate frame with negligible misalignment (**Fig. 1e**). We describe the implementation of RAEM and demonstrate its utility across several biological applications.

## Results

### Implementing the RAEM pipeline

RAEM aims to physically revisit and image specified regions selected within a previously acquired digital volume with nanometer-scale accuracy and resolution. This is enabled by non-destructive serial-section collection methods, including automated tape-collecting ultramicrotomy (ATUM) and direct pickup onto silicon wafers ^**8, 9, 10, 11**^, which mount sections on a stable substrate and thereby establish a consistent, permanent physical coordinate frame for targeting tissue across imaging sessions.

RAEM achieves accurate revisiting by linking the physical sample to a corresponding digital volumetric overview through a coordinate map, *f* (**Fig. 2a**). This map serves two functions: it specifies where on the substrate the microscope should capture image tiles, and it supports stitching and alignment by assembling tiles into 2D section images and elastically aligning those sections to the overview volume.

**Figure 2.**
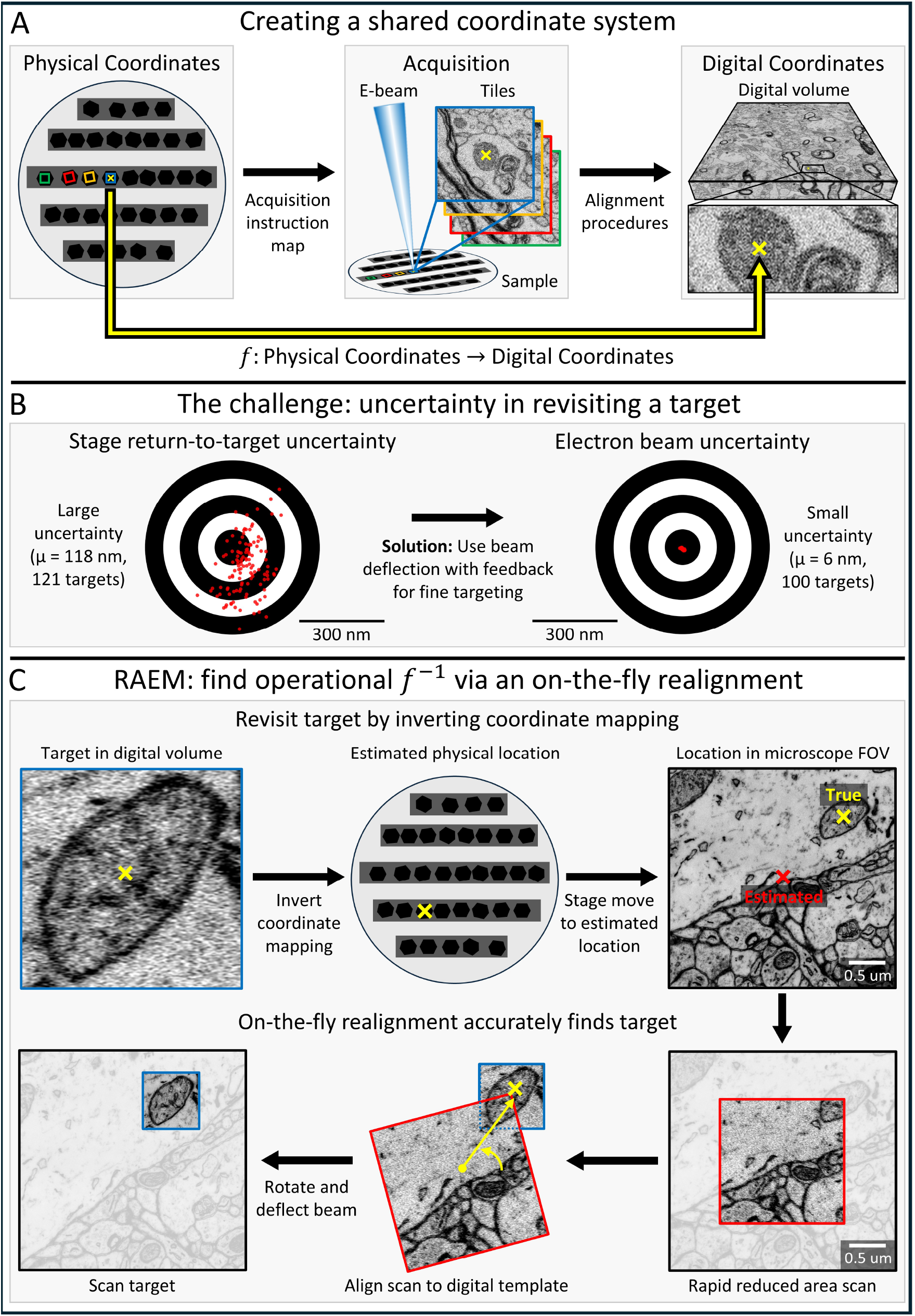
Implementation of the random-access electron microscope (RAEM). **A**. RAEM links physical sample coordinates on the wafer to a stitched and aligned digital volume via a map *f*, composed of an acquisition instruction map (sample → image tiles) and stitching/alignment (image tiles → 3D digital volume). **B**. Revisiting uncertainty. Left: return-to-coordinate uncertainty. A target is imaged, the stage is moved across the wafer to mimic an imaging run, and then returned to the nominal target location for a second image; the misalignment between the two images is measured, yielding ~120 nm on average across 121 targets. Right: electron-beam positioning uncertainty. Two images are acquired at the same location without moving the stage; the misalignment between them is measured, yielding~6 nm on average across 100 targets, comparable to the 4 nm pixel size used during imaging. **C**. RAEM computes an operational inverse map *f* ^*−*1^, which links a digital location to its physical location on the sample, using on-the-fly realignment. A target selected in the digital volume is mapped back to an estimated physical location, revisited by an absolute stage move, then refined by aligning a rapid low resolution reduced area scan to a template from the digital volume. The resulting translation and rotation are applied via beam deflection and scan rotation adjustment to image the target.

Because the map *f* is invertible, a digitally selected pixel can in principle be mapped back to a physical location via *f* ^−1^. However, *f* ^−1^ alone is not sufficient to reliably return to a physical location on a sample, because stage repositioning errors, sample motion, and beam-induced deformation introduce positioning uncertainty.

To characterize this uncertainty, we measured the total return-to-coordinate error across 121 target locations on a sample. At each target, we acquired an image, moved the stage across the 10-cm-scale wafer to mimic section-to-section navigation during volume acquisition, returned to the nominal target location, and acquired a second image. The shift between each image pair yielded an average return-to-coordinate uncertainty of *~*120 nm (**Fig. 2b**, left).

We separately measured electron-beam positioning uncertainty by acquiring two consecutive images of tissue at the same location without moving the stage. The average misalignment was *~*6 nm across 100 targets, comparable to the 4 nm pixel size. This residual reflects both intrinsic beam-positioning uncertainty and any tissue changes between acquisitions, confirming that beam positioning is not the limiting factor in revisiting accuracy.

To compensate for this revisiting uncertainty, RAEM performs on-the-fly realignment (**Fig. 2c**). To image a target selected within the digital overview volume, RAEM first inverts the map *f* to predict the target’s physical location and then revisits that location by stage motion. At the revisited location, RAEM acquires a rapid scan at the center of the field of view and rigidly aligns this scan to a template cropped from the digital volume. The resulting translation and rotation are applied through beam deflection and scan rotation adjustment, correcting the remaining offset with nanometer-scale accuracy.

By achieving nanometer-scale accuracy in revisiting and imaging, RAEM acquires and aligns new image data into the original volumetric overview with negligible misalignment. In the following sections, we demonstrate these revisit-image-fuse capabilities across a range of applications.

### RAEM permits hierarchical and iterative imaging of subregions of interest

In many ssEM applications, structures of interest occupy only a small fraction of the tissue volume. In *C. elegans* connectomics, for example, the entire body must be sectioned, but the nervous system occupies only *~*1% of body volume ^**12, 13, 14, 15, 16**^. Much of the remaining tissue, including muscle and epidermis, need not be imaged at synapse-level resolution and could instead be omitted or acquired at lower resolution. A pipeline that selectively targets the neuropil-rich ganglia and nerve cords would remove a major bottleneck in whole-animal nematode connectomics.

RAEM enables hierarchical imaging of a nervous system embedded within an intact animal. Every serial section is first imaged at low resolution and aligned to generate a volumetric overview, within which regions of interest can then be selected for targeted imaging at higher resolution. This process can be repeated iteratively, with successive acquisitions at spatial scales appropriate to different structures. Because all imaging passes are registered within a common coordinate system, higher resolution data can be integrated seamlessly into the original overview volume.

We demonstrate this workflow in the nervous system of the nematode *P. pacificus*, acquiring the whole animal at low resolution, the head at intermediate resolution, and the brain (nerve ring) at the highest resolution (**Fig. 3a**). Enlarged views of the intermediate and high resolution data (**Figs. 3b, 3c**) show both the improved image quality and the seamless integration of targeted high resolution acquisitions into the surrounding lower resolution context.

**Figure 3.**
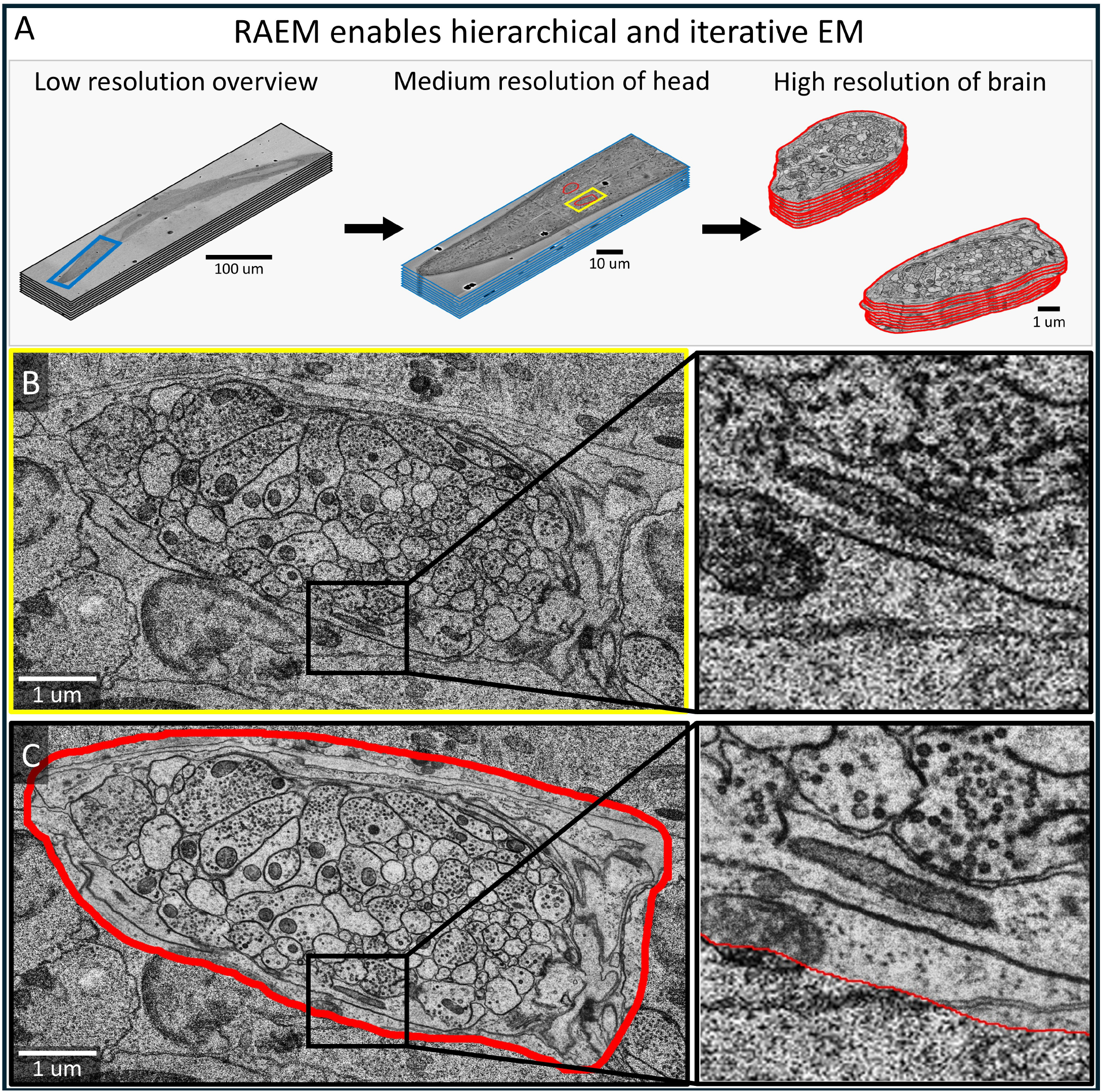
RAEM enables hierarchical, iterative imaging of arbitrary subregions with nanometer-scale accuracy. **A**. Multi-resolution imaging of the nematode *P. pacificus*: a low resolution scan of *N* = 30 serial sections of the whole animal (128 nm, 800 ns), a medium resolution scan of the head (8 nm, 800 ns), and a high resolution scan of the brain (nerve ring, 2 nm, 3000 ns). **B**. Zoom-in of the yellow region in A. **C**. High resolution imaging of the selected subregion, fused back into the medium resolution volume. A magnified view highlights the improved image quality and the precise, seam-free alignment of the high resolution acquisition to the original volume.

An iterative, hierarchical assignment of spatial resolution based on local properties allows imaging time to be allocated efficiently. Regions of sparse neuropil, where neurites are well separated and primarily need to be traced along fibers, can be imaged at lower resolution. In contrast, regions of dense neuropil, where neurites are tightly packed and susceptible to tracing errors, can be targeted for higher resolution acquisition. In this way, RAEM aligns beam time with the demands of downstream reconstruction, transforming EM acquisition from a uniform sampling strategy into a targeted one that adapts to the biological sample.

### RAEM enables targeted, high resolution imaging of mitochondria at scale

Many cell biological studies aim to characterize specific organelles throughout tissue, such as mitochondria. However, ssEM has been difficult to scale to the selective analysis of organelles across large tissue volumes or many samples. Because mitochondria occupy only a small fraction of most cells and tissues, and are small and irregularly distributed, targeted acquisition remains challenging. As a result, traditional ssEM workflows image tissue volumes uniformly at the highest resolution required to resolve functionally important ultrastructural features such as mitochondrial cristae, at the cost of substantial time spent imaging surrounding context ^**17**^.

RAEM makes mitochondrial ultrastructure studies scalable by selectively allocating high resolution imaging time to mito-chondria. To demonstrate this, we applied RAEM to a section of human cortex from the previously published H01 dataset (**Fig. 4a**, left) ^**1**^. We used the pretrained segmentation model MitoNet to detect mitochondria at scale (**Fig. 4a**, middle) ^**18**^. RAEM then revisited and imaged these mitochondria at higher resolution, yielding more than one million mitochondria-centered high resolution 2D images (**Fig. 4a**, right). Because rescans are performed section by section, and because crista ultrastructure can be evaluated in individual cross-sections, we analyzed mitochondria at the single-section level ^**17**^. These targeted scans resolve internal ultrastructural detail, including cristae, that was not visible in the original dataset (**Figs. 4b, 4c**). RAEM-mediated high resolution scanning of human cortex recovers information about mitochondria that was not discernible from the original dataset alone. We extracted mitochondrial morphology and texture features separately from the original low resolution images and the high resolution scans, and then performed principal component analysis (PCA) on each feature set (**Fig. 4d, Extended Data Fig. 1**). Morphological features included area, perimeter, circularity, aspect ratio, solidity, and eccentricity, whereas texture features included intensity coefficient of variation (CV), intensity skewness, intensity kurtosis, edge density, and gray-level co-occurrence matrix (GLCM) descriptors such as contrast, homogeneity, energy, and correlation. Because some features depended strongly on sectioning geometry or imaging conditions, we excluded area, perimeter, intensity CV, and edge density from clustering.

**Figure 4.**
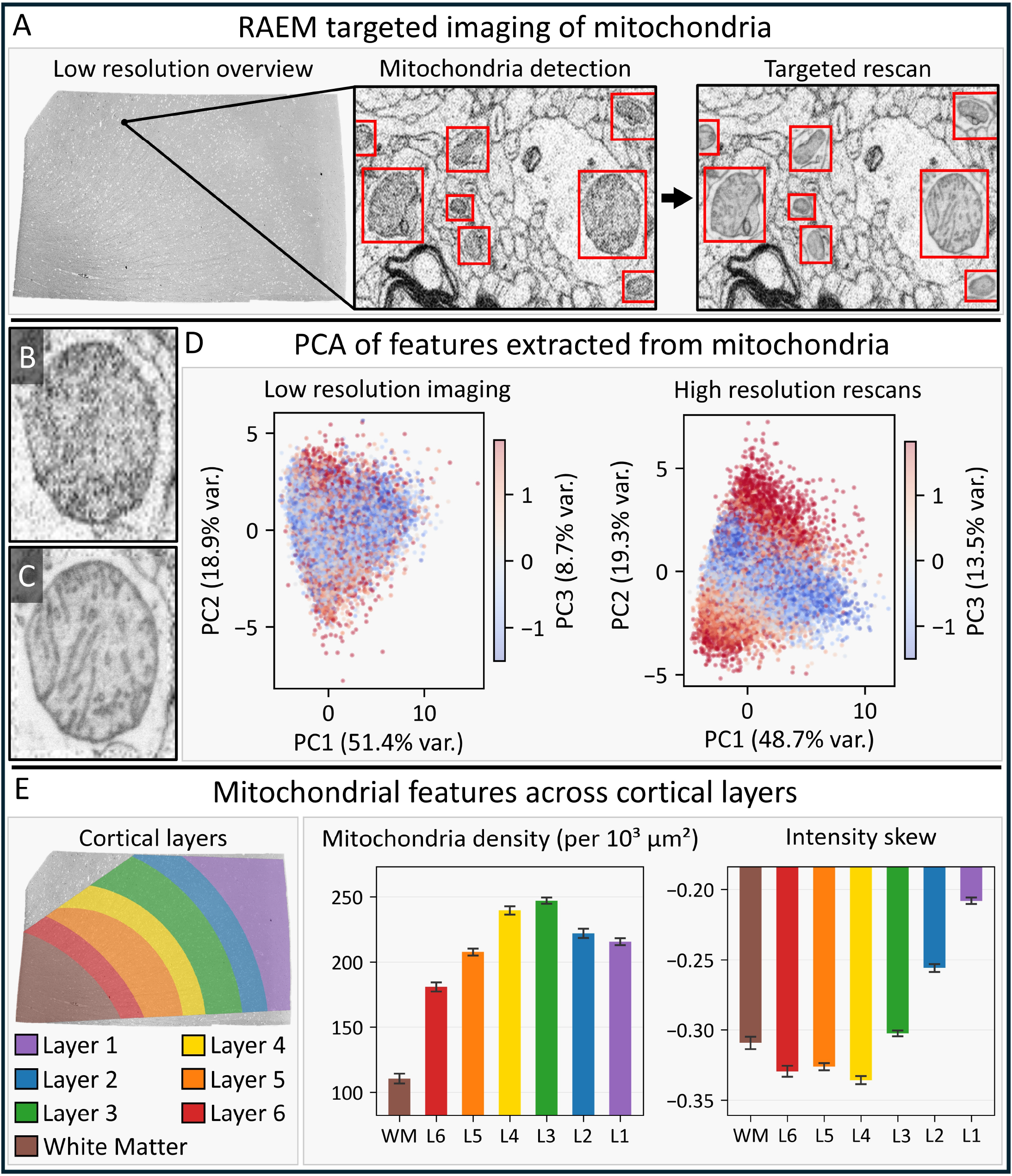
Targeted imaging of over one million mitochondria across human cortex. **A**. Data acquisition pipeline: a low resolution overview (16 nm, 400 ns) of a section of human cortex (H01) is acquired, a neural network (MitoNet) then detects mitochondria, and RAEM rescans each at high resolution (4 nm, 800 ns), yielding *N* = 1,028,096 mitochondria scans after filtering for artifacts. **B**. Example mitochondrion at low resolution. **C**. Same mitochondrion in B at high resolution, resolving cristae ultrastructure. **D**. PCA of morphology and texture features extracted from mitochondria at low resolution (left) and high resolution (right). At high resolution, a texture-dominated axis (PC3) separates cleanly from shape-dominated components, revealing internal ultrastructural variation inaccessible at low resolution. **E**. Cortical layer boundaries overlaid on the H01 section (left). Mitochondrial density (middle) and intensity skewness (right) across cortical layers (density: Kruskal-Wallis *H* = 5,276, *p <* 10^*−*300^, *η*^2^ = 0.33; skewness: *H* = 21,996, *p <* 10^*−*300^, *η*^2^ = 0.024). Mean ± 3×SEM; bars colored by layer.

The first two principal components are dominated by shape features and explain similar amounts of variance at low and high resolution (68% and 70%, respectively), indicating that coarse morphology is captured in both scans (**Extended Data Fig. 1**). In contrast, the third principal component differs strongly between resolutions and reflects texture rather than shape. At high resolution, PC3 explains 13.5% of the variance and loads almost exclusively on intensity-distribution and GLCM texture features (**Extended Data Fig. 1**). It thereby defines a distinct axis of variation, separable from the shape-dominated space defined by PC1 and PC2 (**Fig. 4d**, right).

At low resolution, PC3 explains only 8.7% of the variance, remains entangled with shape (**Extended Data Fig. 1**), and yields a diffuse distribution in PC space (**Fig. 4d**, left), suggesting that internal texture is not reliably resolved. Together, these results show that targeted high resolution scanning uncovers a texture-dominated axis of mitochondrial variation that is inaccessible at lower resolution.

We next asked whether mitochondrial abundance and appearance vary systematically across cortical depth. We transferred cortical layer boundaries from the published H01 dataset onto our section by landmark-based affine registration and assigned each mitochondrion to layers 1–6 or to white matter according to position (**Fig. 4e**, left) ^**1**^.

Mitochondrial density varied strongly across cortical layers (**Fig. 4e**, middle; Kruskal-Wallis *H* = 5,276, *p <* 10^−300^, *η*^2^ = 0.33). Density was highest in layer 3, at 247±0.8 mitochondria per 1000 *µ*m^2^ (mean±SEM across tiles, *n* = 3,569), and lowest in white matter, at 111 ± 1.3 mitochondria per 1000 *µ*m^2^ (*n* = 1,557), corresponding to a 2.2-fold difference. Across cortical depth, density increased from white matter into the deep cortical layers, peaked in layers 3–4, and then declined toward the superficial layers (**Fig. 4e**, middle). Thus, maximal mitochondrial density localizes to the same middle cortical layers in which oxidative metabolism is highest, consistent with greater energy demand in regions of dense synaptic connectivity ^**19**^.

We next examined the texture of mitochondrial internal ultrastructure. The intensity skewness of individual mitochondria differed across cortical layers (**Fig. 4e**, right; Kruskal-Wallis *H* = 21,996, *p <* 10^−300^, *η*^2^ = 0.024, indicating a small but reliable effect). Skewness was most negative in layer 4 (−0.336 ± 0.001; mean ± SEM, *n* = 139,967) and became progressively less negative in the superficial layers, reaching −0.208±0.001 in layer 1 (*n* = 158,335). In EM images, more negative skewness indicates that the intensity distribution within a mitochondrion is shifted toward electron-dense signal. In our data, this signal primarily reflects crista membranes within the organelle ^**17**^. These results suggest that mitochondria are most cristae-dense in layer 4 and exhibit progressively less cristae-dense ultrastructure toward the cortical surface. Other morphology and texture features were statistically significant (Kruskal-Wallis *p <* 10^−50^ for all) but showed small effect sizes across layers (**Extended Data Figs. 2a, 3**).

We next asked whether mitochondrial structure is uniform or instead organized into discrete subtypes across cortical layers. We used two complementary analyses. First, k-means clustering (*k* = 2–15) on the first six principal components yielded uniformly low silhouette scores (**Extended Data Fig. 2b**), with a maximum of 0.37 at *k* = 2. Second, t-SNE embeddings across perplexities of 10–200 produced a single connected cloud, with no fragmentation into discrete groups (**Extended Data Fig. 2c**). Together, these results suggest that mitochondrial morphology varies along a continuum rather than partitioning into discrete subtypes. We nevertheless used k-means partitions (*k* = 5) as a practical way to divide this continuum and examine how these groups localize across cortical layers (see Supplementary Information).

We note that padded mitochondria-targeted scans in the human cortex section covered only approximately 7% of the tissue area. Acquiring this *~*6.9 mm^2^ section by conventional uniform scanning at 4 nm pixel size and 800 ns dwell time would have required approximately 95 hours of beam time, whereas our targeted acquisition required only approximately 8.5 hours, representing an 11-fold reduction. RAEM therefore makes it feasible to acquire and compare high resolution ultrastructural measurements of organelles such as mitochondria across large tissue areas and diverse tissue types. More broadly, this pipeline should enable rapid comparative studies of mitochondrial ultrastructure across tissues and experimental conditions.

### RAEM enables vesicle-resolved synapse imaging with targeted 3D scans

We next tested RAEM on synaptic boutons, another class of biologically important structures whose key ultrastructural features require higher resolution than the surrounding tissue context to resolve. We acquired and aligned a low resolution volumetric overview of human cortex spanning 165 ×140 ×2 *µ*m^3^ (**Fig. 5a**, left). From this digital volume, we segmented 20 synaptic boutons with padding (**Fig. 5a**, middle) and then acquired each bouton in 3D at higher resolution (**Fig. 5a**, right).

**Figure 5.**
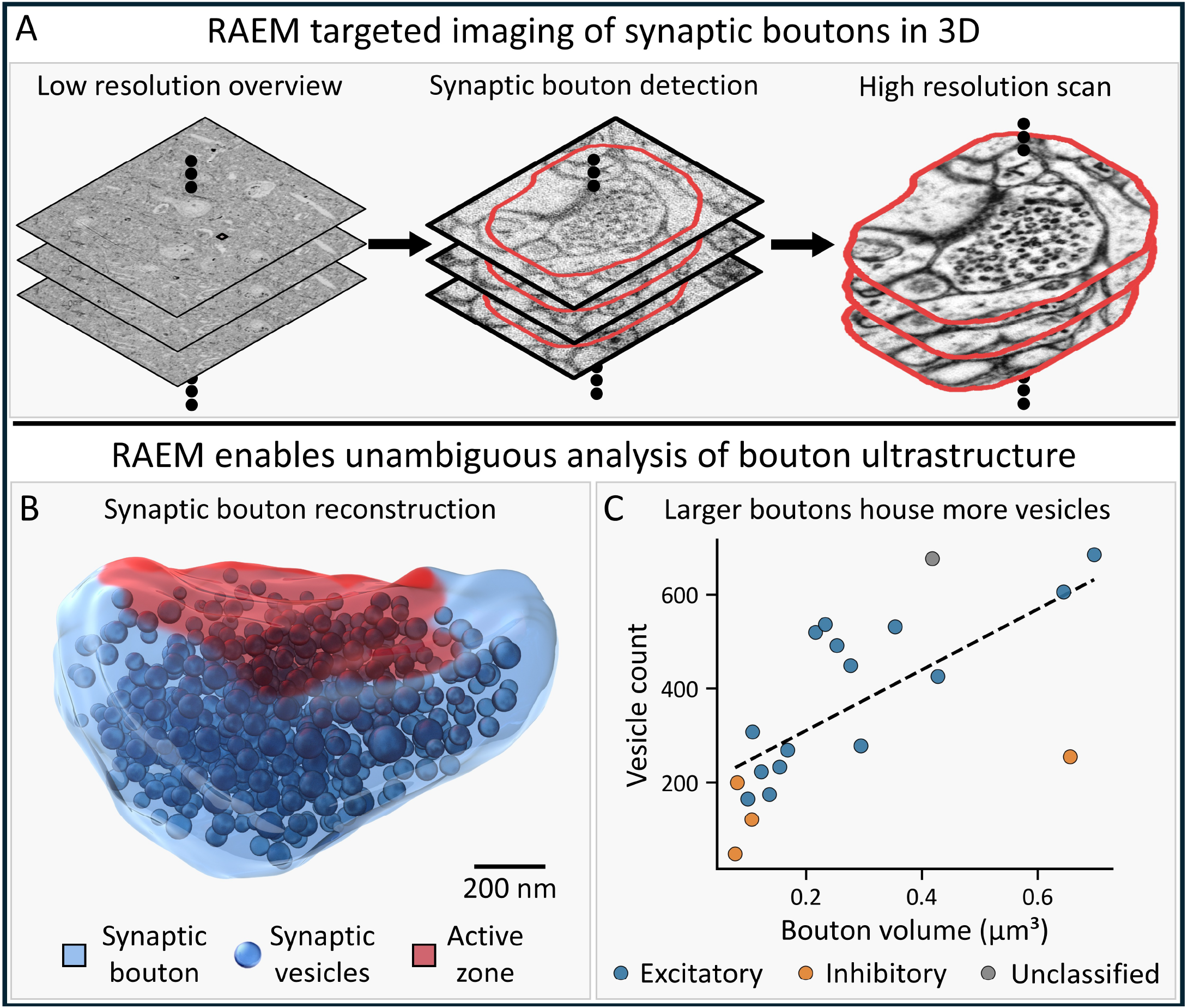
Targeted imaging of synaptic boutons in 3D in human cortex. **A**. Data acquisition pipeline: a low resolution overview volume (8 nm, 200 ns, 165 ×140 ×2 *µm*^3^, *N* = 60 sections) is acquired and aligned (left), synaptic boutons are identified in 3D with padding (middle), and each is scanned at high resolution (4 nm, 1200 ns, right). **B**. 3D reconstruction of a bouton. **C**. Vesicle count vs. bouton volume for 20 boutons, colored by synapse type (excitatory, inhibitory, or unclassified). Larger boutons house more vesicles (Pearson *r* = 0.67, *p* = 0.001, *n* = 20 boutons).

In the initial low resolution overview, bouton ultrastructure was largely unresolved. Features such as synaptic vesicles were indistinct and could not be identified reliably (**Extended Data Fig. 4a**). Comparison with the same bouton in the published H01 multibeam SEM dataset showed better contrast, but vesicle ultrastructure remained ambiguous (**Extended Data Fig. 4b**). By contrast, RAEM high resolution targeted scanning resolved individual vesicles clearly enough to support reliable detection and counting (**Extended Data Fig. 4c**).

RAEM targeted scans also resolved vesicle ultrastructure that was not discernible in the original multibeam SEM dataset. For example, we could reliably identify clear-core vesicle morphology across serial sections. Thus, RAEM targeted scanning improved both vesicle detection and three-dimensional morphological classification (**Extended Data Fig. 4d**).

These rescans enabled quantitative analysis of the 20 boutons. We segmented each bouton in 3D, measured its volume, and counted its vesicles (**Fig. 5b, Extended Data Fig. 5**). To assign synapse type, we coregistered the rescanned boutons to the published H01 volume and used its existing automated synapse-sign classifications. Of the 20 boutons, 15 were classified as excitatory, 4 as inhibitory, and 1 remained unclassified. Bouton volumes ranged from 0.077 to 0.697 *µ*m^3^, and vesicle counts ranged from 47 to 685. These quantities were positively correlated (**Fig. 5c**; Pearson *r* = 0.67, *p* = 0.001, *n* = 20), consistent with larger boutons containing more synaptic vesicles. We also examined vesicle density by synapse type and found a trend toward lower vesicle density in inhibitory boutons, although the sample size was insufficient to establish statistical significance (**Extended Data Fig. 4e**). Together, these analyses illustrate new kinds of quantitative ultrastructural measurements that RAEM now makes possible.

### RAEM enables revisiting of previously acquired volumes

RAEM can be implemented as an end-to-end pipeline in which the same single-beam SEM acquires both the low resolution overview and the high resolution targeted scans. Alternatively, overview images from pre-existing datasets acquired on a different microscope can be registered into the RAEM coordinate system and used to define targets for acquisition on the RAEM single-beam SEM.

We tested this alternative workflow using the H01 dataset, which was originally acquired by multibeam SEM. For a given serial section in H01 (**Fig. 6a**, left), we acquired a low reso-lution overview of the same physical section using our single-beam SEM and affine aligned the published H01 section to this overview (**Fig. 6a**, middle). Affine alignment alone brought the multibeam SEM and single-beam SEM datasets into agreement to within *~* 30 *µ*m across 33 sampled locations. To correct the remaining mismatch, we estimated a local deformation field that captures elastic differences between the multibeam dataset and the RAEM single-beam overview (**Fig. 6a**, right). We computed this field by acquiring medium resolution single-beam images at grid points across the section and aligning each image to its predicted location in the published H01 dataset, as determined by the affine transform (**Extended Data Fig. 6**). After applying this deformation field, residual offsets fell within the range correctable by RAEM’s on-the-fly beam realignment. End-to-end rescanning at three locations spanning millimeters of tissue confirmed residual offsets below 4 *µ*m, well within RAEM’s on-the-fly correction range.

**Figure 6.**
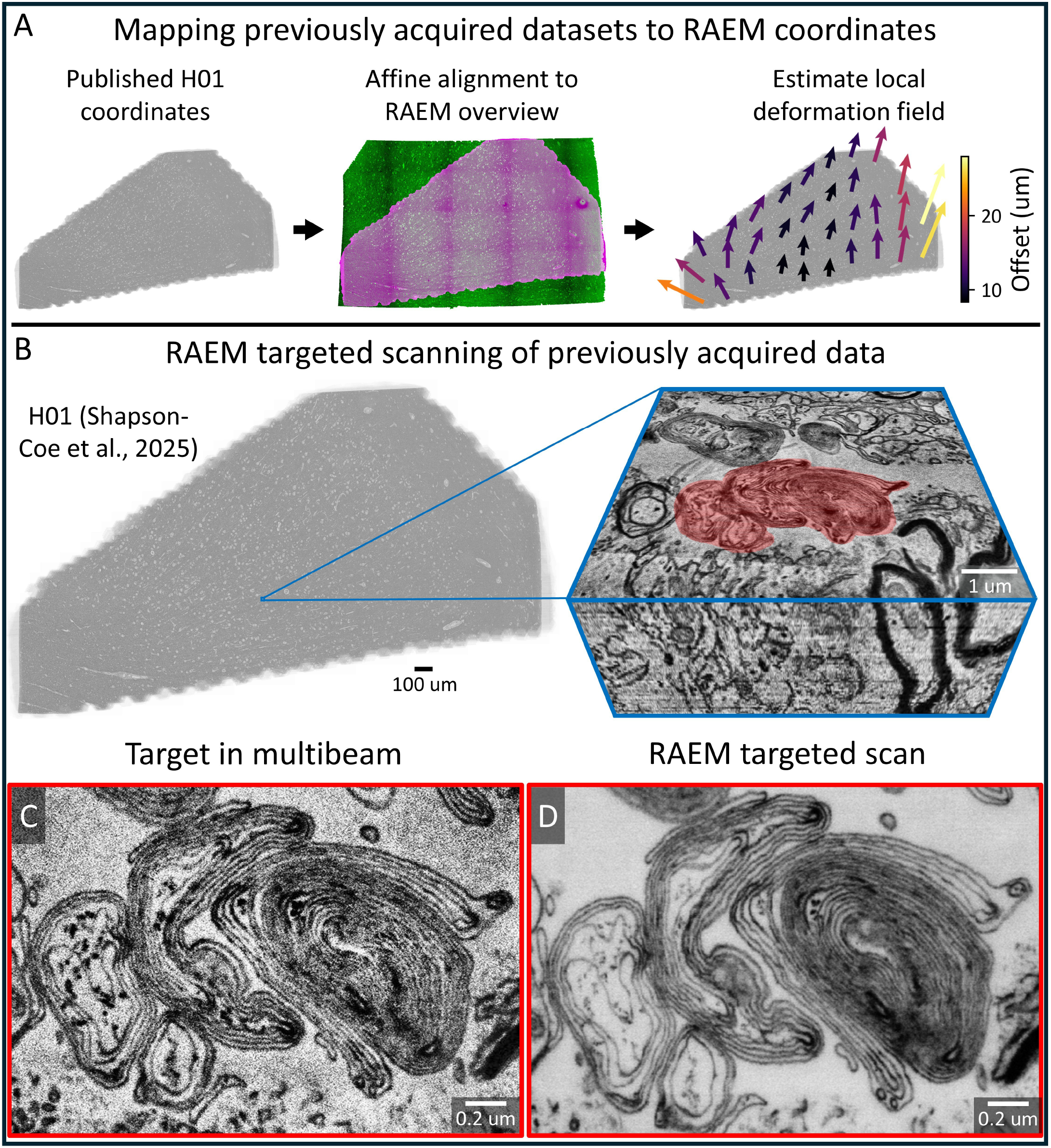
Revisiting previously acquired volumes via cross-modality registration. **A**. Mapping a published dataset to RAEM coordinates: a section from the published H01 multibeam volume (left) is affine aligned to a RAEM single-beam overview (middle; magenta: published, green: RAEM overview), and a local deformation field corrects residual misalignments (right). **B**. Overview of the published H01 section with a target of interest highlighted and shown in 3D. **C**. The same target in the published H01 multibeam volume. **D**. RAEM single-beam rescan of the same target.

With the H01 section registered to the RAEM coordinate frame, RAEM can then revisit and image arbitrary targets from the published dataset directly on the single-beam SEM (**Figs. 6b, 6c, 6d, Extended Data Fig. 7**).

This proof of concept shows that volumes acquired on other microscopes can be made compatible with RAEM revisiting. More broadly, it extends the utility of RAEM by enabling hybrid workflows in which large tissue volumes are first screened rapidly by multibeam SEM, and selected regions are then revisited at higher resolution on a single-beam SEM. In the examples presented here, RAEM rescanning on a single-beam SEM with backscattered electron detection yielded higher quality ultra-structural images than the original multibeam dataset, enabling more informative follow-up imaging of selected targets.

## Discussion

RAEM’s revisiting capabilities extend beyond EM-only workflows. Any imaging modality that can be registered to the RAEM coordinate frame could, in principle, provide targets for subsequent EM acquisition. For example, a light microscopy volume could be aligned to a low resolution EM overview, allowing fluorescently labeled structures to be revisited and imaged at high resolution by EM. This suggests a path toward correlative workflows in which lower resolution light microscopy modalities guide ultrastructural follow-up by RAEM.

By establishing a consistent, persistent coordinate frame, RAEM has the potential to change how EM samples are used. Once the mapping between sample and digital volume has been established, any region of tissue can be revisited and imaged within a shared spatial framework, with each new acquisition incorporated into the existing volume. In this view, EM samples are no longer consumed by a single analysis, but can instead be revisited in iterative studies. A volume first acquired for one purpose, such as tracing neurons, could later be revisited to analyze synapses, organelles, or other features as new questions emerge.

Thus, RAEM enables a collaborative model of ultrastructural biology. A mapped tissue volume could function as a shared reference resource to which different groups contribute targeted acquisitions addressing distinct biological questions. Repeated electron exposure could in principle degrade tissue, but resin-embedded samples tolerate substantial dose before visible damage, and in our experiments we observed no detectable degradation across multiple targeted imaging passes. RAEM also points toward new opportunities for computationally guided image acquisition. At present, SmartEM operates section by section and therefore uses only 2D information to identify regions that require higher resolution imaging to correct segmentation errors. A next step would be to extend such approaches into 3D, allowing volumetric context to identify truly error-prone regions more selectively. In practice, many apparent ambiguities in 2D may be resolvable from adjacent sections without additional rescanning. Incorporating 3D context could further reduce the amount of rescanning required and push SmartEM toward greater efficiency.

A practical limitation of the current pipeline is that it operates one wafer at a time. Revisiting regions across wafers requires an additional alignment procedure, as illustrated here through registration to the H01 dataset. Future extensions of the pipeline could make such cross-wafer and cross-dataset registration increasingly routine. This limitation may also be mitigated by modern section-collection methods such as magnetic collection, which routinely place hundreds to thousands of sections on a single wafer and should provide sufficient volumetric extent for many targeted imaging applications ^**10, 11**^.

The idea of using image-based feedback to revisit locations on serial-section substrates was explored by WaferMapper, which cross-correlated images with references and applied corrections through additional stage moves ^**20**^. RAEM instead applies corrections through beam deflection and scan rotation, achieving nanometer-scale targeting without additional stage moves.

## Acknowledgements

Research reported in this paper was supported by the NIH BRAIN Initiative under grant no. U01NS132158 (awarded to A.D.T.S. and J.W.L.). The authors acknowledge BossDB.org for providing data storage and access services that support the sharing of the datasets associated with this publication (grant no. R24MH114785) ^**21**^.

## Author Contributions Statement

Conceptualization: I.S.C., Y.M., A.D.T.S. and J.W.L. Methodology and software: I.S.C. led the overall implementation. M.P. and I.S.C. performed the bouton analysis. Y.W. and D.B. provided technical guidance. R.S. assisted with sample handling and microscope operation. Supervision and writing: the project was supervised by Y.M., A.D.T.S. and J.W.L. The manuscript was written by I.S.C., Y.M., A.D.T.S. and J.W.L. with input from all authors.

## Extended Data

**Extended Data Figure. 1.**
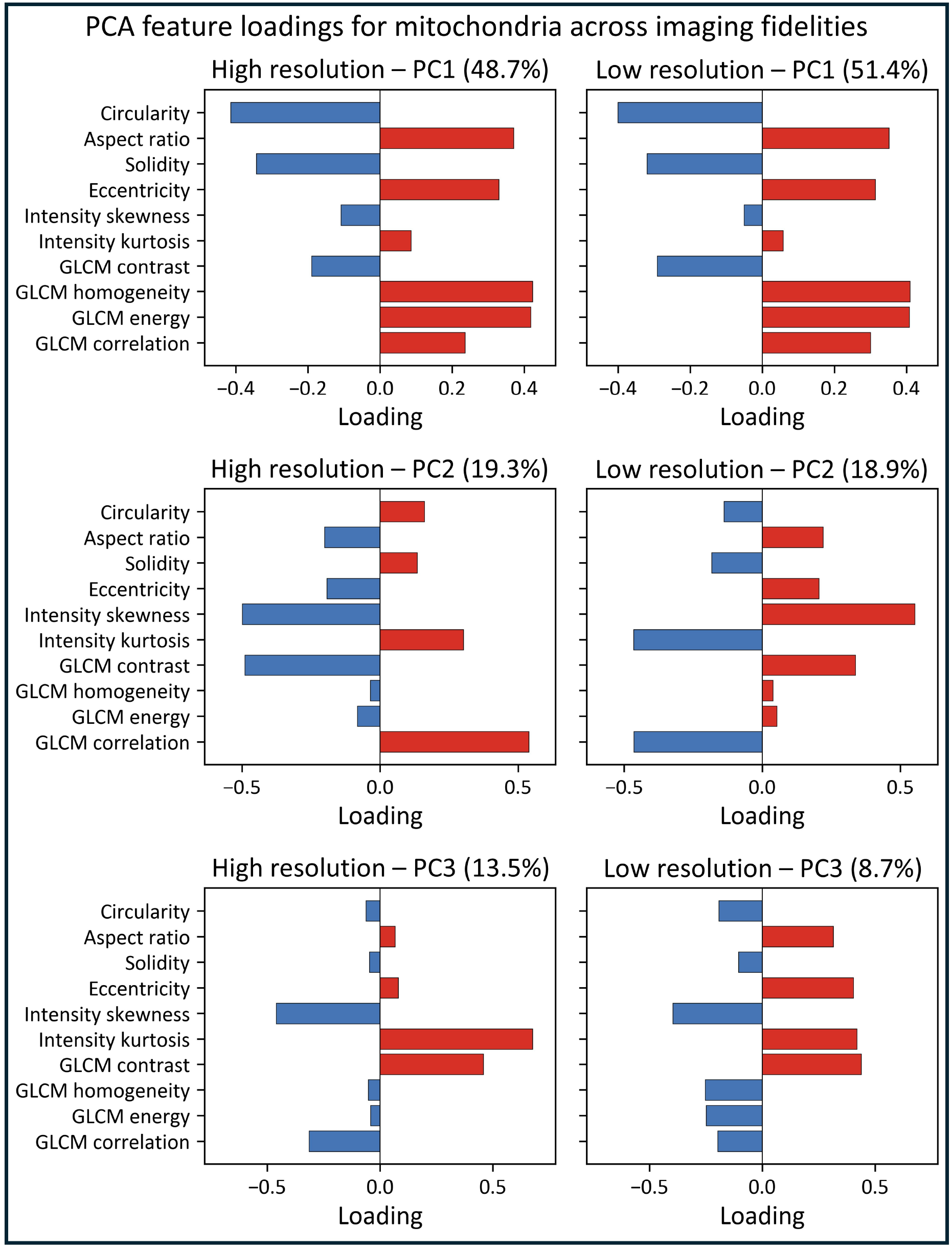
PCA feature loadings for mitochondria across imaging resolutions. Loading weights of morphology and texture features on the first three principal components, computed separately from low resolution (16 nm, 400 ns) and high resolution (4 nm, 800 ns) mitochondria images. At both resolutions, PC1 and PC2 are dominated by shape features. PC3 differs: at high resolution, it loads almost exclusively on texture features such as intensity distribution and GLCM descriptors, whereas at low resolution, it remains entangled with shape features such as eccentricity.

**Extended Data Figure. 2.**
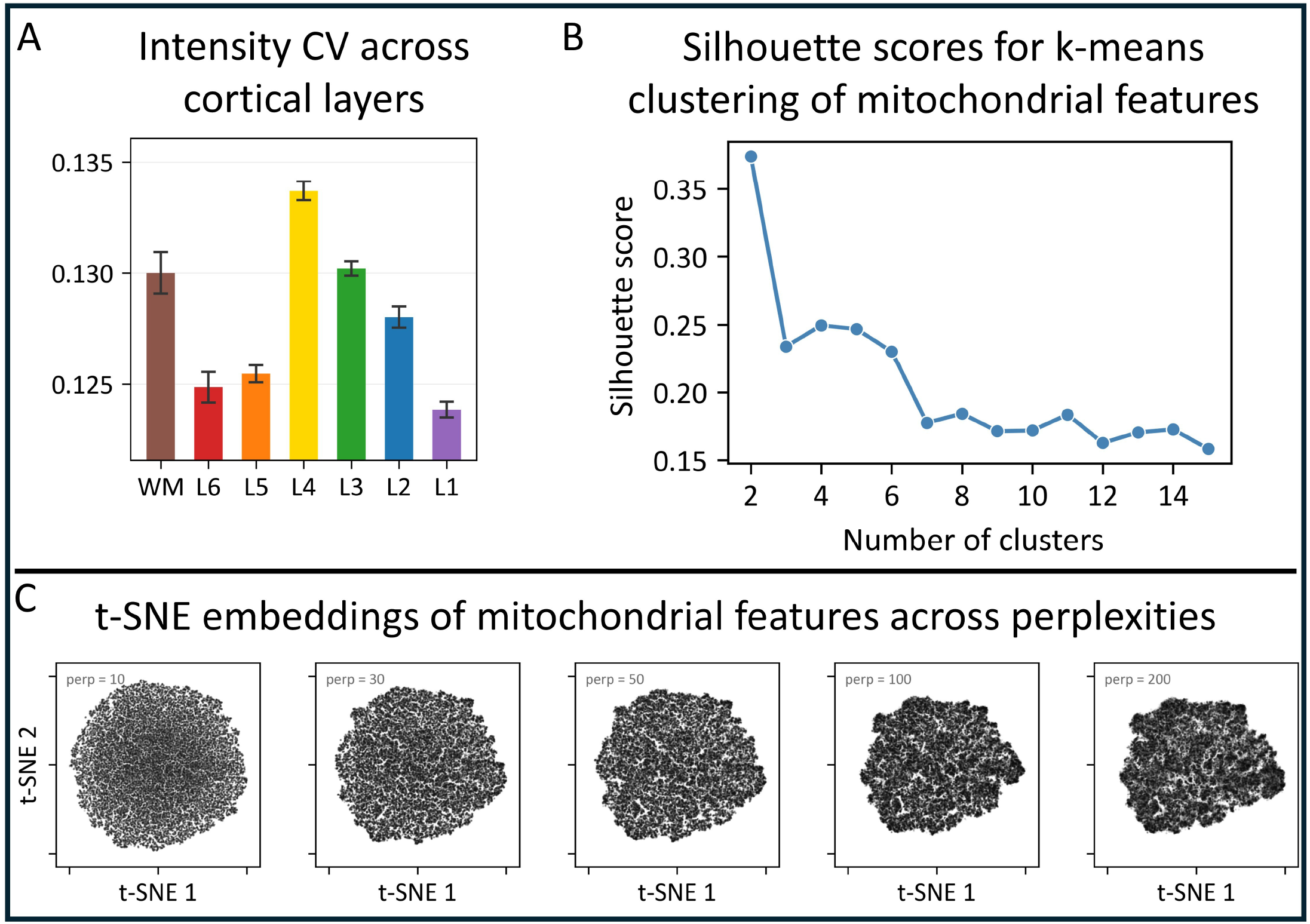
Additional mitochondrial analyses. **A**. Intensity CV peaks in layer 4 (Kruskal-Wallis *H* = 10,834, *p <* 10^*−*300^, *η*^2^ = 0.012). **B**. Silhouette scores for k-means clustering (*k* = 2–15) on the first six principal components, yielding uniformly low scores (maximum 0.37 at *k* = 2). **C**. t-SNE embeddings across perplexities 10–200, producing a single connected cloud with no fragmentation into discrete groups. B and C suggest that mitochondrial morphology varies along a continuum rather than falling into discrete subtypes.

**Extended Data Figure. 3.**
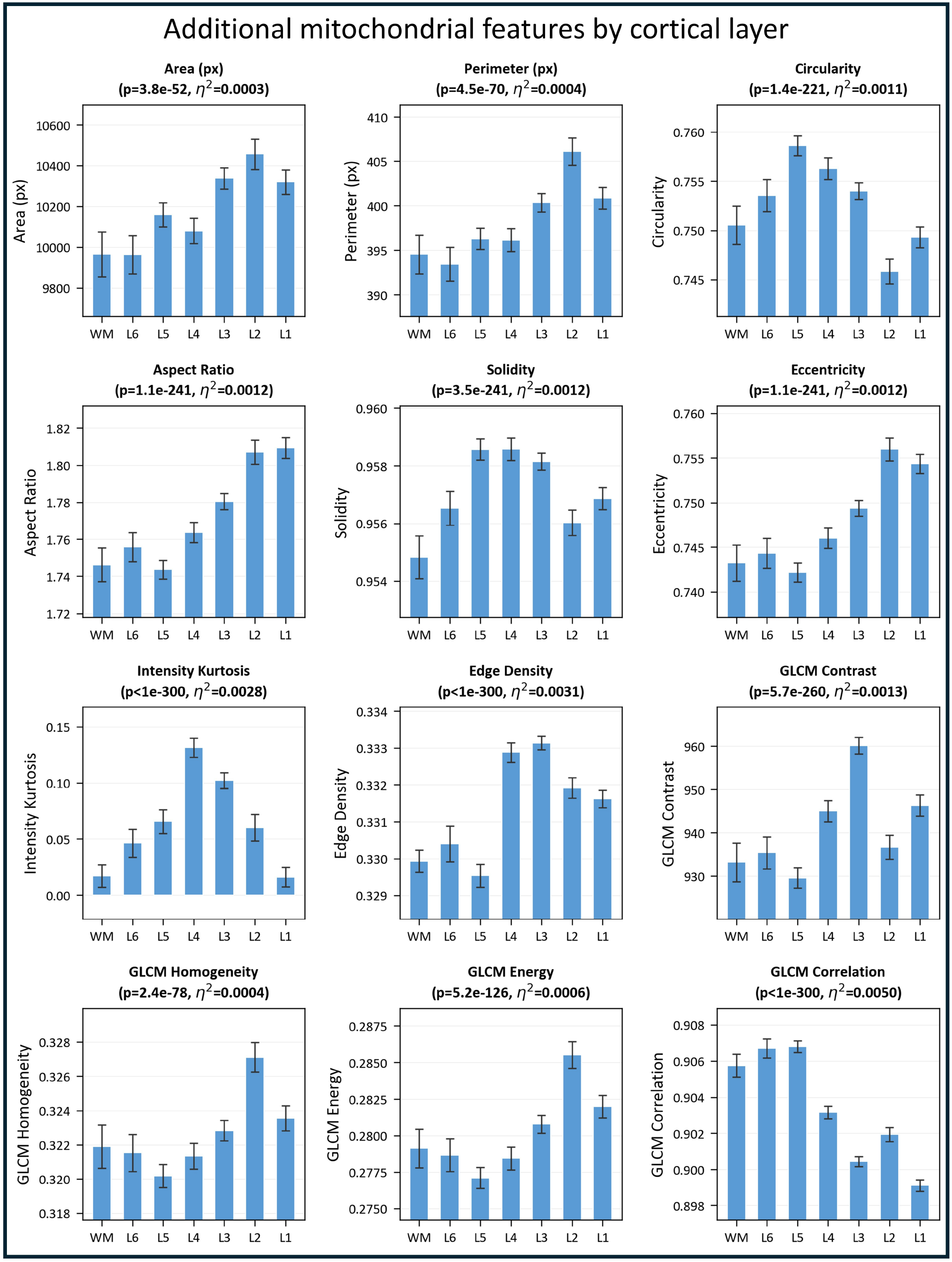
Mitochondrial morphology and texture features across cortical layers. All features show statistically significant variation across layers (Kruskal-Wallis *p <* 10^*−*50^ for all) but with small effect sizes. Bars depict mean ± 3×SEM.

**Extended Data Figure. 4.**
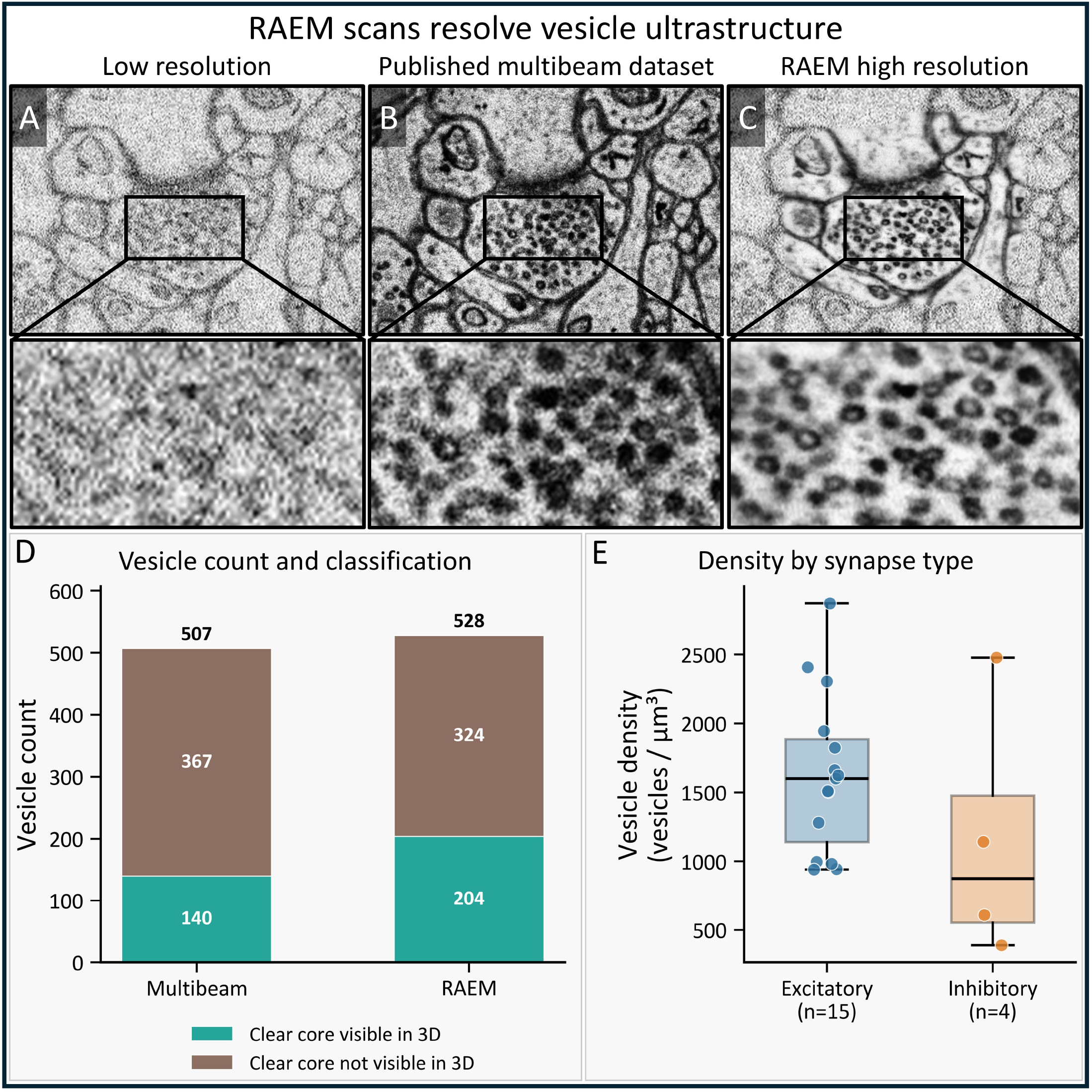
Targeted high resolution imaging with RAEM improves vesicle detection and classification. **A**. Low resolution RAEM scan of a synaptic bouton, acquired to identify targets of interest. **B**. The same bouton as it appears in the published H01 multibeam SEM volume. **C**. High resolution targeted RAEM scan of the same bouton, resolving individual vesicle ultrastructure including clear-core morphology. **D**. Vesicle counts and 3D classification for this bouton imaged by both modalities. RAEM detected 4% more vesicles overall (528 vs. 507) and, critically, identified 46% more vesicles whose clear core was visible in serial sections (204 vs. 140). The remaining vesicles lacked a discernible clear core across serial sections in either modality. **E**. Vesicle density (vesicles/*µ*m^3^) across 20 boutons segmented from the RAEM volume, grouped by synapse type. Inhibitory boutons appear to pack fewer vesicles per unit volume than excitatory boutons (median 876 vs. 1601 vesicles/*µ*m^3^), though this difference did not reach significance (Mann–Whitney *U, p* = 0.262), likely due to the small inhibitory sample size (n=4).

**Extended Data Figure. 5.**
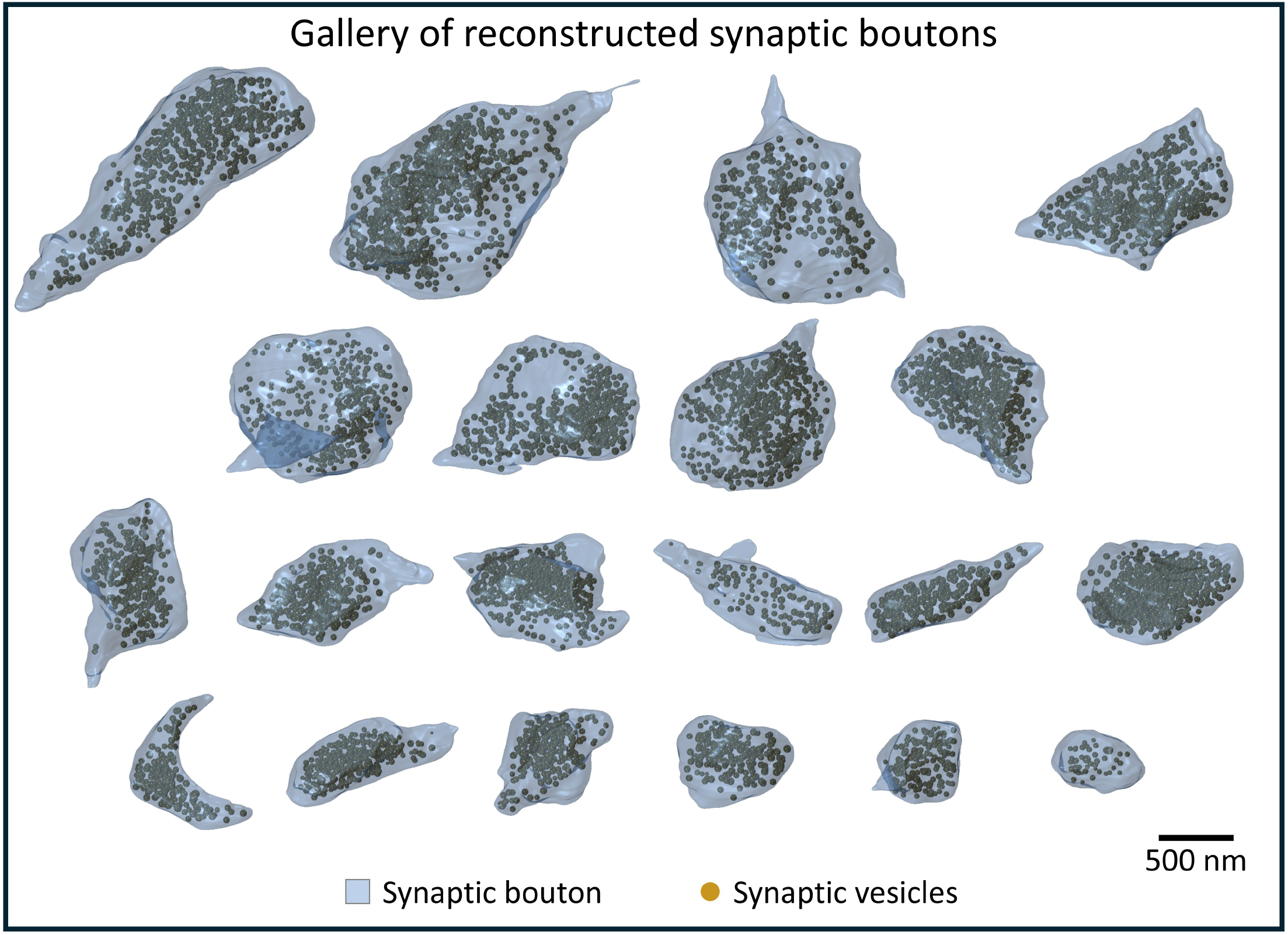
Gallery of reconstructed synaptic boutons. 3D reconstructions of 20 synaptic boutons (blue, semi-transparent) with synaptic vesicles (gold) from the H01 dataset, rendered at the same scale and arranged by volume (largest to smallest).

**Extended Data Figure. 6.**
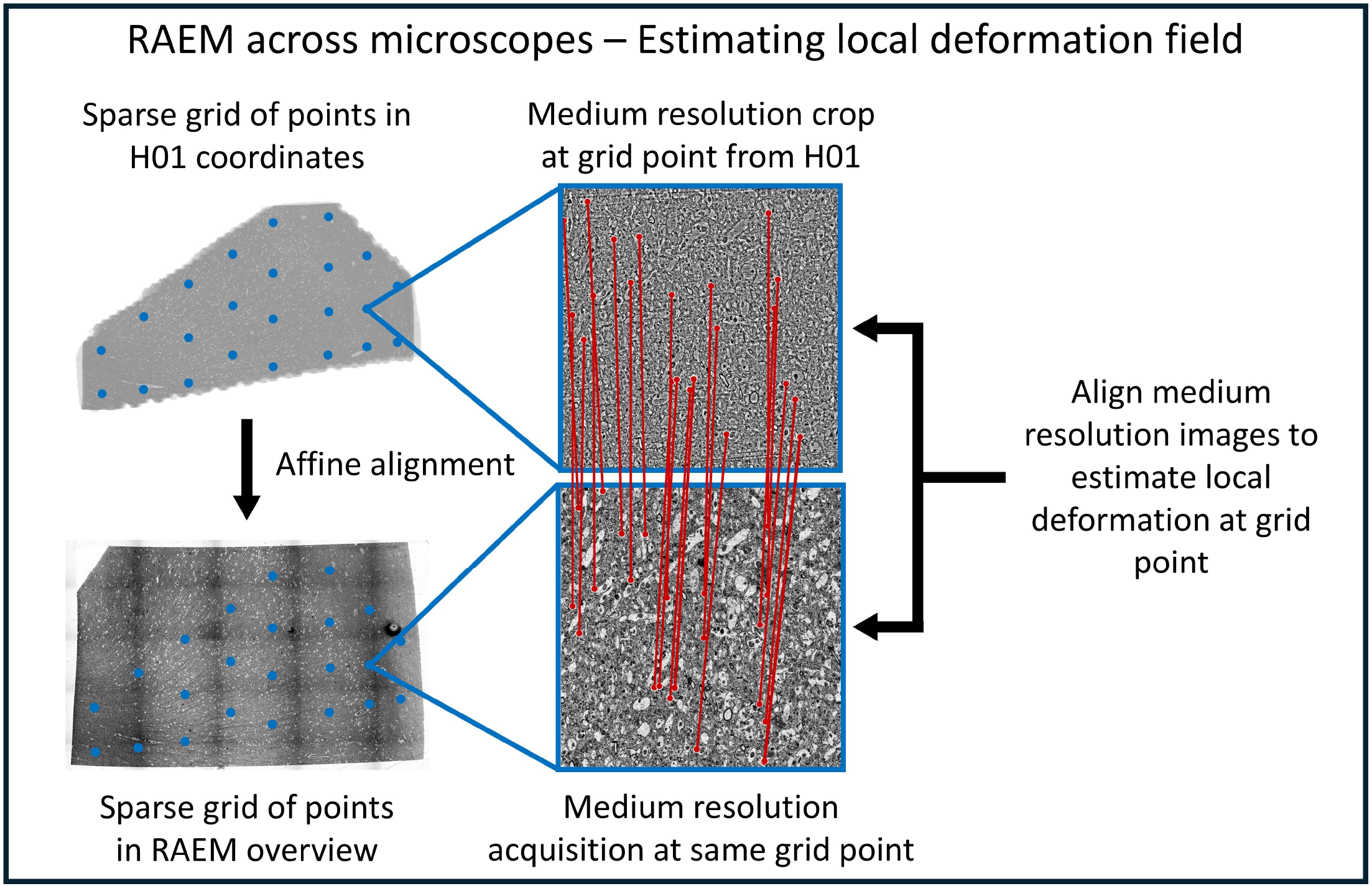
Estimating the local deformation field between published H01 and RAEM coordinates. A sparse grid of locations is selected in the published H01 volume and mapped to RAEM overview coordinates via an affine alignment between the low resolution views of the two datasets (left). At each grid point, a medium resolution image is acquired by single-beam SEM and aligned to a template cropped from the corresponding location in the published multibeam volume (middle). Red lines indicate SIFT feature correspondences between the two modalities. Repeating this procedure across all grid points yields a spatially resolved estimate of the local deformation field, which corrects residual distortions not captured by the global affine transform.

**Extended Data Figure. 7.**
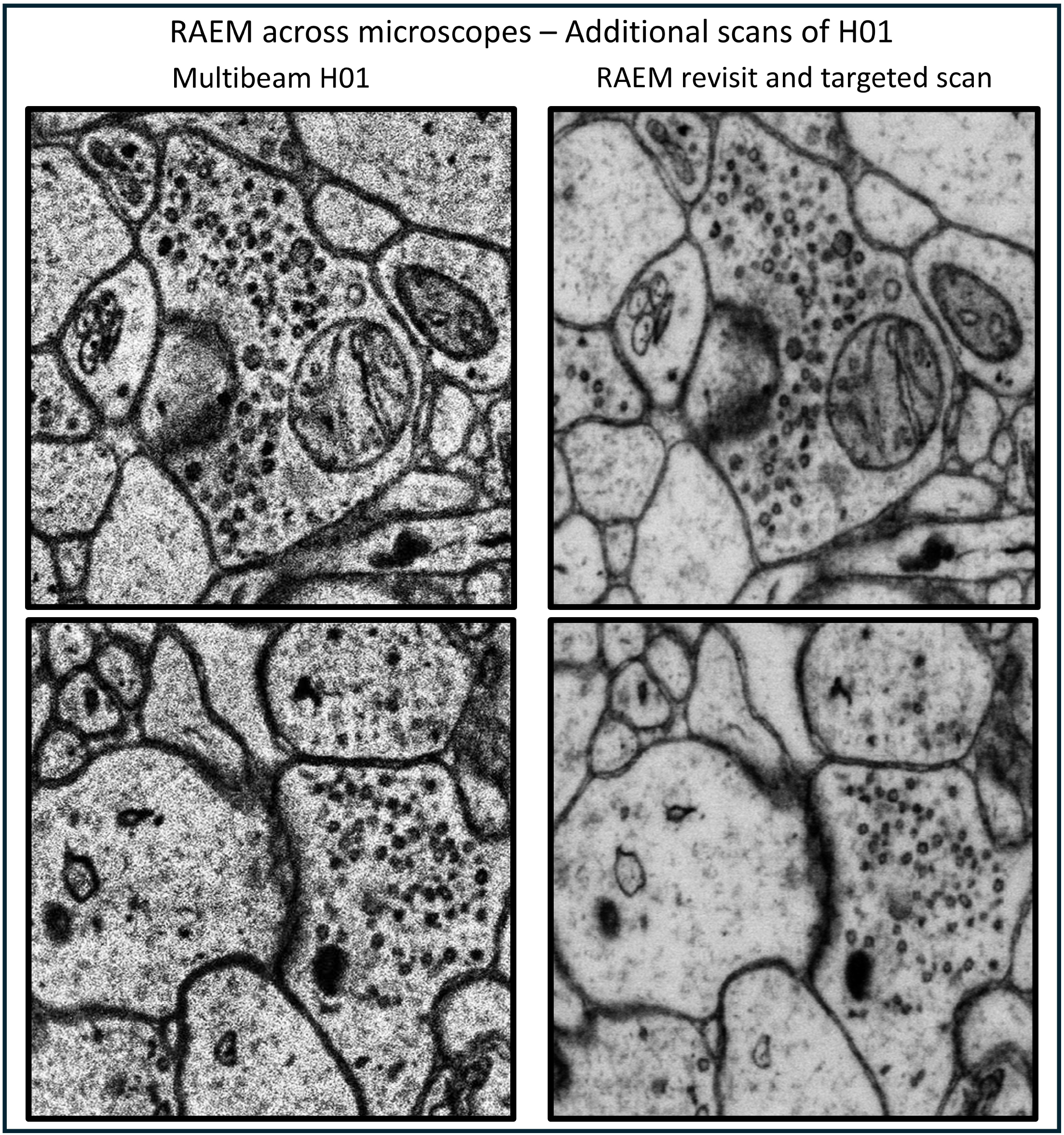
Additional cross-modality images of the H01 dataset. Each pair shows the same region in the published H01 dataset (left) and the corresponding RAEM single-beam targeted scan (right).

## Methods

### Imaging procedure

#### Microscope hardware and imaging parameters

All imaging was performed on a Thermo Fisher Verios 5 HP scanning electron microscope equipped with a concentric backscattered electron (CBS) detector at 6.5 mm working distance and 4000 V stage bias. Landing energy was 7 kV for all experiments. Beam current was 1.6 nA for nematode tissue and 6.4 nA for human cortex tissue. Serial sections on the wafer were identified and targeted using SEM Navigator, a custom software akin to the previously published WaferMapper ^**20**^, and all microscope control was performed programmatically via the Thermo Fisher AutoScript API (https://www.thermofisher.com/us/en/home/electron-microscopy/products/software-em-3d-vis/autoscript-4-software.html).

#### Sample preparation and section collection

Nematode tissue was prepared via high-pressure freezing and freeze substitution as described previously in refs. ^**14, 22**^. Thirty-nanometer sections were cut with a Leica EM UC6 ultramicrotome and collected onto kapton tape using an automated tape-collecting ultramicrotome (ATUM). Tape was mounted on silicon wafers, and sections were post-stained with uranyl acetate and lead citrate ^**8**^. Human cortex tissue was previously published as the H01 dataset; sample preparation and section collection are described in ref. ^**1**^.

#### Stitching and elastic alignment

Overlapping image tiles were stitched into 2D sections and aligned into 3D volumes using FEABAS. Briefly, block matching was used to find correspondences between overlapping tile pairs, and triangular meshes were relaxed to produce seamless montages. For 3D alignment, stitched sections were downsampled to generate thumbnails, which were aligned across serial sections via feature matching. Fine alignment was then performed at higher resolution using template matching, and elastic transformations were computed via mesh relaxation using a sliding-window optimization strategy. Code for stitching and alignment is available at https://github.com/YuelongWu/feabas.

#### Estimating the inverse map *f* ^−1^

To revisit a target selected in the digital volume, RAEM must compute *f* ^−1^, mapping a digital coordinate back to a physical location on the sample. The forward map *f* is composed of two transformations: (1) an acquisition instruction map, which links physical stage coordinates to raw image tiles, and (2) elastic stitching and alignment, which maps raw tiles into the 3D digital volume. To estimate *f* ^−1^, RAEM maps the target’s coordinate in the digital volume back through the inverse of the elastic alignment to a specific raw tile. This is done by identifying the mesh triangle containing the target point via R-tree spatial lookup and applying barycentric interpolation on the mesh vertices to obtain the corresponding coordinate in the raw tile. Since the acquisition instruction map records the physical stage position of each tile, the target’s pixel offset from the center of that raw tile is converted to a physical distance using the known pixel size and added to the tile’s stage position. A rotation correction is applied to account for the scan rotation angle and the inversion between image and stage coordinate systems. The resulting coordinate is a predicted physical stage position on the sample. RAEM then refines this estimate via on-the-fly realignment. We note that when the coordinate is revisited, the beam rotation for this revisit is initialized based on the orientation of the target in the digital volume, rather than the original scan angle, and is refined during on-the-fly realignment.

#### On-the-fly realignment

After the stage moves to the predicted physical location, the positioning is not sufficiently accurate for targeted imaging due to stage repositioning uncertainty. RAEM corrects the remaining offset by acquiring a rapid low resolution preview at the center of the field of view and rigidly aligning it to a template cropped from the digital volume. A partial affine transform is estimated via SIFT feature matching ^**23**^ with RANSAC outlier rejection and reduced to a rigid transform (rotation and translation) via Procrustes alignment. The resulting translation offset is applied via beam deflection to recenter the field of view on the target, and the scanning rotation is adjusted accordingly.

#### Targeted imaging

Targeted imaging uses the microscope’s patterning engine to scan only selected pixels, as previously described in ref. ^**7**^. A binary bitmap mask, where nonzero pixels mark regions to scan, is generated from targets selected or detected in the overview volume. At each field of view, after on-the-fly realignment computes the in-plane offset and rotation, the bitmap is loaded into the patterning engine and positioned using the realignment offset and rotation. The patterning engine steers the beam exclusively to pixels within the bitmap, resulting in a sparse scan that covers only the targets of interest.

#### Targeting strategies for targeted scanning

Three approaches determine which locations to visit during targeted scanning. The first places each target within a single field of view, suitable when targets fit within one field of view and the number of targets per section is small, as in the nematode and bouton experiments. When targets are numerous or larger than a single field of view, the section is covered with an overlapping tile grid and only tiles that overlap with at least one target are acquired; targets that span multiple tiles are stitched afterward, as in the mitochondria experiment. An alternative approach for numerous small targets groups them into cliques that can be scanned together within a single field of view, without requiring post-acquisition stitching. There is a tradeoff between the latter two strategies depending on target density and distribution.

#### Fusion to digital volume and optional post-processing

Because the on-the-fly realignment already registers each targeted scan to the digital volume, targeted acquisitions can be fused directly into the overview volume. Small residual offsets due to local elastic warping in the digital volume and imprecision in beam steering result in minor misalignments between the targeted scan and the overview. As an optional refinement, each targeted scan is affine aligned to the corresponding region of the digital volume for pixel-perfect fusion. To remove edge artifacts, targeted scans are eroded by 1 pixel at the boundary before fusion, as described in ref. ^**7**^.

#### Measuring revisiting uncertainty

For both return-to-coordinate and beam positioning measurements, images were acquired at 4 nm pixel size, 1200 ns dwell time (2048 *×* 1768 pixels) on nematode tissue. For return-to-coordinate uncertainty, images were acquired at 121 locations, the stage was moved to mimic a full imaging run, and images were reacquired at the same locations. For beam positioning uncertainty, two images were acquired consecutively at each of 100 locations without moving the stage; the measured misalignment reflects both intrinsic beam uncertainty and any changes in tissue shape between acquisitions. In both experiments, misalignments between image pairs were measured via SIFT feature matching with RANSAC-based rigid registration, and all alignment transforms were validated by manual inspection.

### Application: Targeted imaging of the nematode nervous system

#### Nematode hierarchical and iterative imaging

Thirty serial sections of the nematode *P. pacificus* were imaged at three resolutions. The whole animal was imaged at 128 nm pixel size, 800 ns dwell time (1024 *×* 884 pixel tiles, 10% overlap), tiled into a mosaic, and aligned in 3D with cross-correlations and SIFT respectively. The head region was identified in the overview and imaged at 8 nm pixel size, 800 ns dwell time (6144 *×* 6144 pixel tiles, 10% overlap). The nerve ring was identified in the medium resolution volume and reimaged at 2 nm pixel size, 3000 ns dwell time. The medium resolution pass was stitched and aligned using FEABAS. The high resolution pass was fused into the digital volume via RAEM’s fusion capabilities.

### Application: Targeted imaging of mitochondria in human cortex

#### Overview acquisition

A single section of human cortex from the H01 dataset (section 1113) was imaged at 16 nm pixel size, 400 ns dwell time (6144 *×* 4096 pixel tiles, 8% overlap). The section area was ~6.9 mm^2^, acquired in a serpentine scan pattern with autofocus sites spaced at least 300 µm apart. Auto-contrast/brightness calibration was performed at the start of each row.

#### Mitochondria segmentation

Mitochondria were segmented from the overview mosaic using MitoNet via the empanada framework in semantic mode ^**18**^. Each tile was preprocessed with contrast-limited adaptive histogram equalization (CLAHE) and Gaussian blur, and padded with neighboring tiles to provide boundary context. After inference, padding was discarded and cross-tile instance merging was performed using a union-find algorithm on edge strips at tile boundaries. A minimum area filter was applied to remove spurious detections (0.04 *µm*^2^). Manual validation of segmentation quality on five randomly sampled tiles (359 detected mitochondria) yielded a precision of 0.99 and recall of 0.83.

#### Feature extraction

Fourteen morphometric features were computed for each segmented mitochondrion from the high resolution (4 nm) images. Six shape features were measured: area, perimeter, circularity, aspect ratio, solidity, and eccentricity. Three intensity features were computed from pixel values within each mitochondrial mask: coefficient of variation, skewness, and kurtosis. Five texture features were extracted: edge density and four gray-level co-occurrence matrix (GLCM) descriptors–contrast, homogeneity, energy, and correlation. GLCM descriptors were computed at a distance of 1 pixel across four angles (0°, 45°, 90°, 135°) with 256 gray levels, and the four directional values were averaged for each property. Pixel intensities within each mitochondrial mask were rescaled to the full [0, 255] range before GLCM computation. For dimensionality reduction and clustering, 10 features were used, excluding area, perimeter, intensity coefficient of variation, and edge density. Area and perimeter were excluded because they depend on sectioning geometry, and intensity coefficient of variation and edge density were excluded for sensitivity to imaging conditions.

#### Per-tile normalization and destriping

Texture and intensity features were z-score normalized within each tile to remove tile-to-tile variations in imaging conditions. Shape features were not normalized as they are invariant to imaging conditions. We observed systematic horizontal banding in some texture features across tile rows, caused by auto-contrast/brightness variations during acquisition. We accounted for this by subtracting the row mean and adding back the global mean for each feature, removing the banding without distorting biological gradients.

#### Cortical layer assignment

Cortical layer boundaries were transferred from the published H01 dataset onto our section via landmark-based affine registration. The H01 EM image and layer segmentation were downloaded at mip9 resolution, and a full affine transform was computed from manually placed correspondences between the H01 image and the RAEM overview. The resulting layer mask assigns labels for layers 1-6 and white matter. An exclusion mask was manually drawn to remove regions with sectioning artifacts. Each mitochondrion was assigned to a cortical layer based on its centroid position in the layer mask. The transform was computed from 8 manually placed landmark pairs; because layer boundaries span hundreds of micrometers, residual registration error (on the order of tens of micrometers) is unlikely to affect layer assignments except near layer transitions.

#### Statistical analysis

Between-layer differences were assessed using the Kruskal-Wallis H-test for each feature across seven cortical layers. Effect sizes were quantified using eta-squared. Mitochondrial density was computed per tile (4096 × 4096 pixels), with each tile assigned to a cortical layer based on the layer mask at its center.

### Application: Vesicle-resolved imaging of synaptic boutons

#### Volume acquisition and targeted scanning

A volume of human cortex spanning 165 × 140 × 2 *µm*^3^ (60 serial sections) was acquired at 8 nm pixel size, 200 ns dwell time. Tiles were stitched and elastically aligned using FEABAS. Twenty synaptic boutons were manually segmented in 3D from the aligned volume with contextual padding around each target. Each bouton was scanned at 4 nm pixel size, 1200 ns dwell time.

#### Vesicle counting and volume estimation

Vesicles and bouton volumes were manually segmented in 3D by an expert annotator using VAST ^**24**^. Bouton volumes were estimated by voxel counting from the segmentation masks.

#### Synapse type classification

Each bouton was identified in the published H01 volume, and synapse type (excitatory or inhibitory) was determined using the automated synapse sign classifications from the published dataset ^**1**^.

#### 3D rendering

Bouton surfaces were reconstructed from segmentation masks using a custom Python script. Masks were resampled to isotropic voxels, morphologically closed and hole-filled, Gaussian smoothed (*σ* = 10 nm), and isosurfaced via marching cubes. Meshes were further smoothed with iterative Laplacian relaxation. Each vesicle was rendered as a sphere with diameter estimated from its cross-sectional area, assuming a circular profile.

### Application: Revisiting a previously published dataset

#### Cross-modality registration

To align the published H01 multibeam volume to the RAEM single-beam overview, we acquired a low resolution overview of the same physical section in field-free mode, which permits much larger fields of view suitable for low resolution screening, and computed a global affine transform between the two images via SIFT feature matching with RANSAC. Affine alignment alone brought the two datasets to within ~30 *µm* of alignment. To correct the remaining spatially varying misalignment, we acquired medium resolution images at a sparse grid of positions across the section in immersion mode (immersion mode permits higher resolution imaging at the expense of smaller fields of view) and aligned each to the corresponding region in the published volume. The translational offsets at each grid point define a local deformation field, interpolated via Delaunay triangulation. After applying this correction, residual offsets fell within the range correctable by RAEM’s on-the-fly realignment. Because the overview was acquired in field-free mode and targeted scans use immersion mode, a fixed offset between the two modes was measured and applied during targeted scanning.

## Data Availability

All datasets are available via a BossDB ^**21**^ project page at https://bossdb.org/project/chandok2026.

## Code Availability

All code produced for this study is available on GitHub at https://github.com/nano6626/RAEM.

## Supplementary Information

### RAEM system and workflow

**Supplementary Figure. 1.**
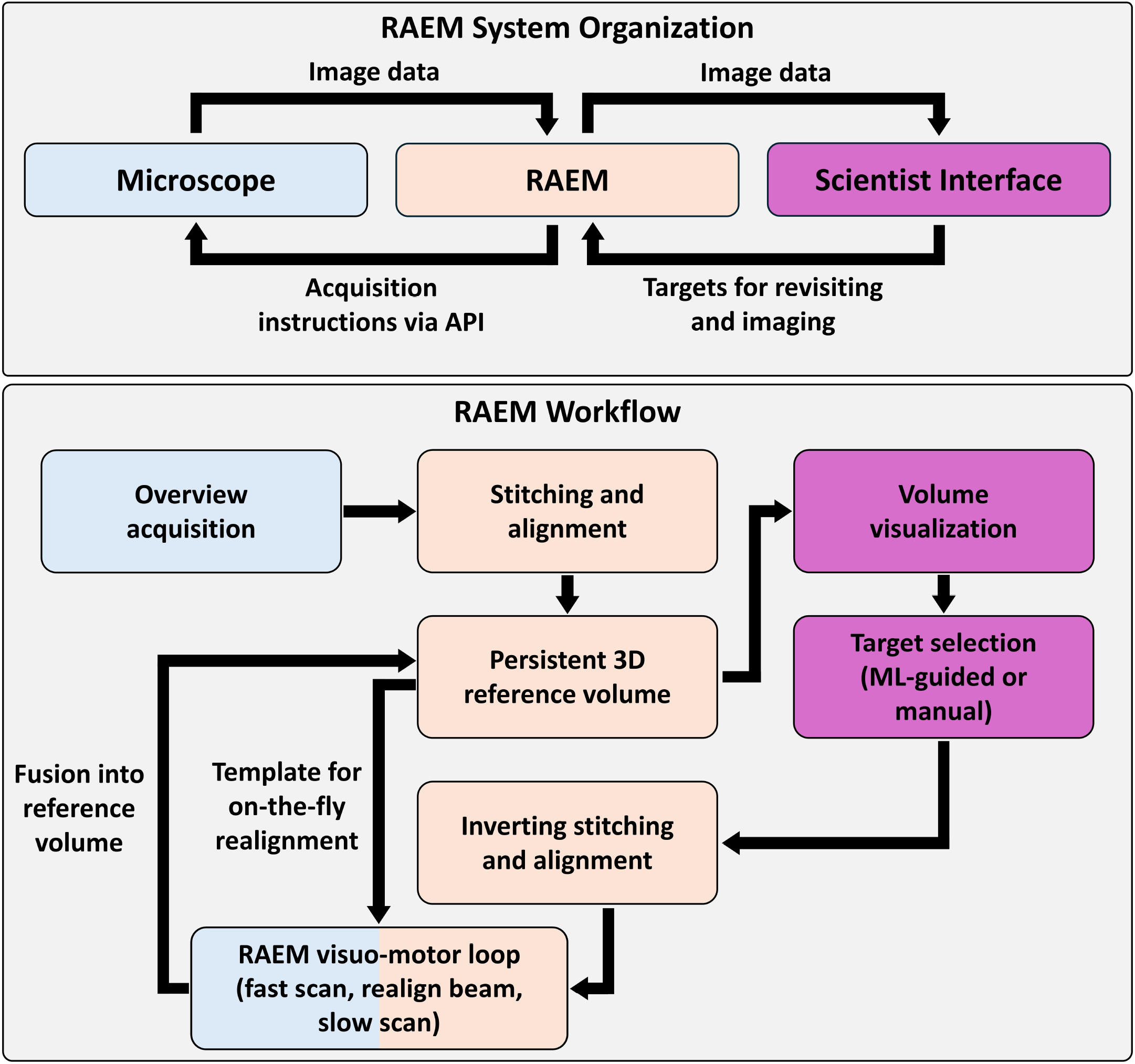
RAEM system organization and workflow. Top: RAEM comprises three modules. The microscope acquires image data and receives acquisition instructions (stage movements, beam parameters) from RAEM via an API. RAEM sends image data to a scientist interface, where a scientist or machine learning (ML) model can visualize the data and select targets for high resolution imaging. Selected targets are sent back to RAEM. Bottom: The RAEM workflow begins with overview acquisition, followed by stitching and alignment to produce a persistent 3D reference volume. This volume is visualized in the scientist interface, where targets are selected manually or by ML. To image a target, RAEM inverts the stitching and alignment to recover physical sample coordinates, then executes a visuo-motor loop: a fast scan is acquired, the beam is realigned on-the-fly using a template from the reference volume, and the target is imaged at high resolution. The resulting targeted scan is fused back into the reference volume. This process can be repeated iteratively for any target in the volume.

### Clustering of mitochondrial features

Silhouette analysis and t-SNE visualization indicate that mitochondrial morphology in this tissue varies along a continuum rather than partitioning into discrete subtypes (**Extended Data Fig. 2b, 2c**). We nevertheless adopted a *k* = 5 partition as a practical means of summarizing the dominant axes of morphological variation. Clustering was performed on 10 features, four describing shape (circularity, aspect ratio, solidity, eccentricity) and six describing texture (intensity skewness, intensity kurtosis, GLCM contrast, homogeneity, energy, and correlation). These features were standardized and projected onto six principal components explaining 96.8% of the variance. Area and perimeter were excluded because they depend on sectioning geometry, and intensity CV and edge density were excluded due to sensitivity to imaging conditions. These four features are reported for completeness but did not contribute to cluster assignments.

The five clusters and their feature profiles are summarized in **Supplementary Figs. 2a, 2b**. Cluster 1 (35.6%) contains compact, round profiles (circularity 0.84) with dense, evenly distributed cristae. Cluster 2 (22.9%) contains moderately elongated profiles (aspect ratio 2.16) with similar internal texture to cluster 1. Cluster 3 (8.0%) is distinguished by its internal texture rather than shape, with strongly negative intensity skewness (−0.96) and elevated kurtosis (1.83) reflecting high internal contrast. Prototypical profiles often contain prominent dark structures within the interior. Cluster 4 (23.5%) consists of small profiles (mean area approximately 0.89 *µm*^2^) with grainy internal texture. Cluster 5 (10.0%) contains highly elongated profiles (aspect ratio 3.02) with irregular, non-elliptical contours (solidity 0.85).

We examined whether the relative proportions of these five classes varied across cortical layers (**Supplementary Fig. 2c**). No clear pattern of cluster composition emerged: the five clusters were represented in broadly similar proportions across layers 1–6 and white matter. This is consistent with the small effect sizes of most clustered features across layers (*η*^2^ *<* 0.03); the layer-associated gradients in these features are too subtle relative to within-layer morphological diversity to produce spatially segregated clusters.

**Supplementary Figure. 2.**
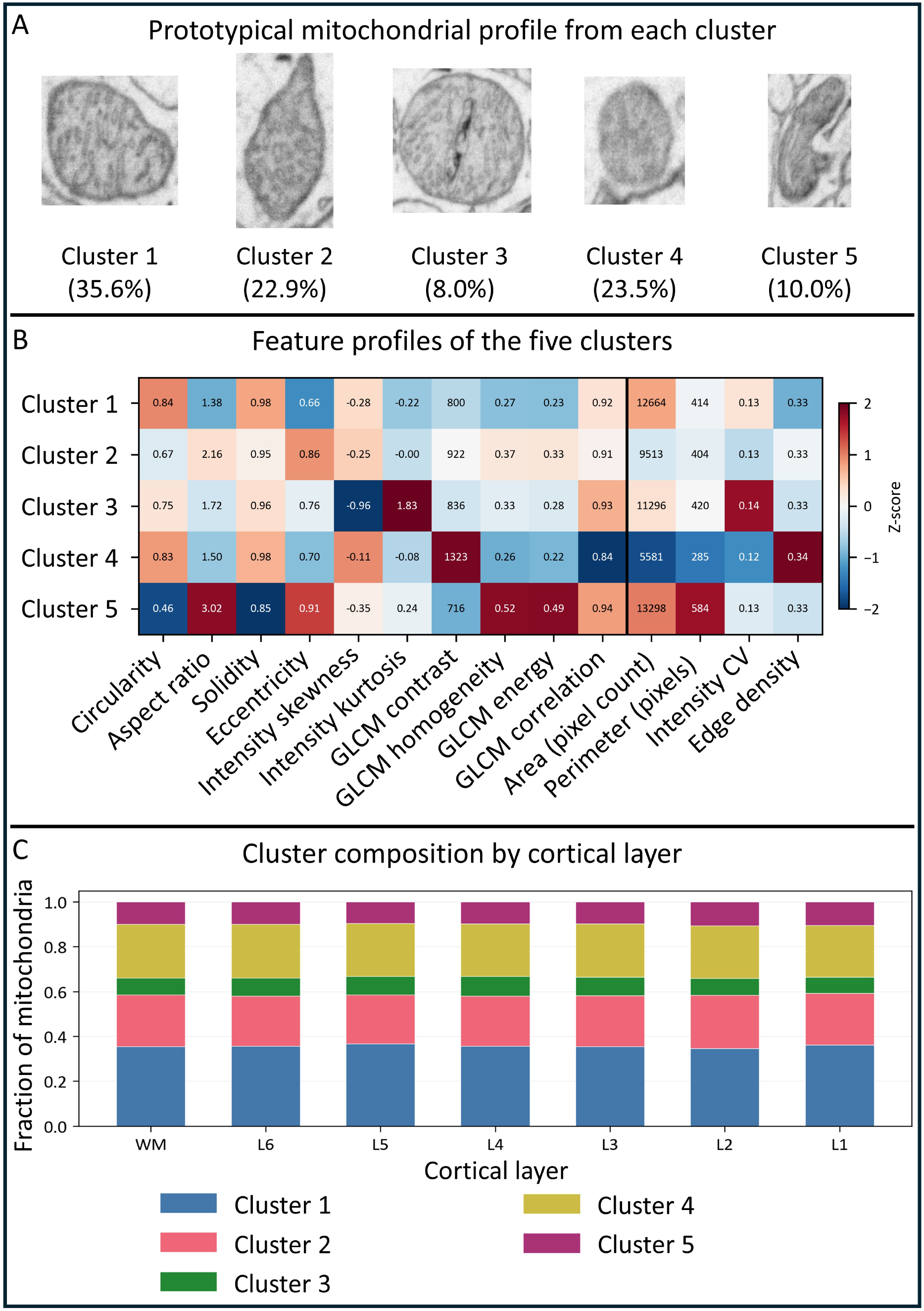
K-means clustering of mitochondrial morphology. **A**. Representative mitochondrial profile from each of the five clusters. **B**. Mean feature values across clusters (z-scored). Colors represent z-scores computed across the five cluster means for each feature independently, placing all features on a comparable scale regardless of their original units. Raw (un-normalized) cluster mean values are annotated within each cell. A vertical line separates the 10 features used for clustering (left) from four additional features (area, perimeter, intensity CV, edge density; right) that were measured but excluded from clustering. **C**. Cluster composition across cortical layers. Stacked bars show the relative proportion of each cluster within each cortical layer and white matter.

